# Distribution and abundance of the West Indian manatee (*Trichechus manatus*) in the Panama Canal

**DOI:** 10.1101/026724

**Authors:** Giselle Muschett, Juliana A. Vianna

## Abstract

The West Indian manatee (*Trichechus manatus*) is threatened throughout its distribution, and is categorized as vulnerable by IUCN (Lefebvre 2001, IUCN 2007). The number of mature individuals is currently estimated at less than 10,000 and is expected to decline at a rate of at least 10% over the next 20 years as a result of both habitat loss and anthropogenic factors (Deutsch *et al*. 2008). While the West Indian manatee is the most widespread of all existing sirenians, some populations are estimated at fewer than 10 individuals (Deutsch *et al*. 2008, Quintana-Rizzo and Reynolds 2010). In addition, in most Central American countries reliable information on manatee distribution and conservation status is missing and there is need to assess these remnant populations to guide future management strategies.

---

In Panama, there has been little recent manatee research (Muschett *et al*. 2009, Lefebvre 2001). There are two known resident manatee populations in the country, one in Bocas del Toro, on the Northern Caribbean Coast, and the other in Lake Gatun, in the Panama Canal Watershed (Fig. 1) (Mou-Sue *et al*. 1990, Lefebvre 2001). The origin of this second population is unclear, and whether manatees in the Chagres River survived the construction of the Panama Canal remains unknown (Mou-Sue *et al*. 1990, MacLaren 1967). However, in 1964 one Amazonian manatee *T. inunguis* from Peru and nine West Indian manatees *T. manatus* from Bocas del Toro were introduced into an enclosure in Lake Gatun as part of an aquatic vegetation control program for the Panama Canal. Some years later these manatees either escaped or were released into the lake (MacLaren 1967). Since then manatee sightings have been common, as have vessel collisions and deaths from underwater detonations for dredging (Schad *et al*. 1981, Hernández 1982). However, an aerial survey carried out years later spotted only one manatee in the lake (Mou-Sue *et al*. 1990). To date the actual number of manatee in Lake Gatun remains undetermined. The annual number of deaths is also unknown and there are unconfirmed reports of hunting by local people (Lefebvre *et al*. 2001).

**Figure 1.**
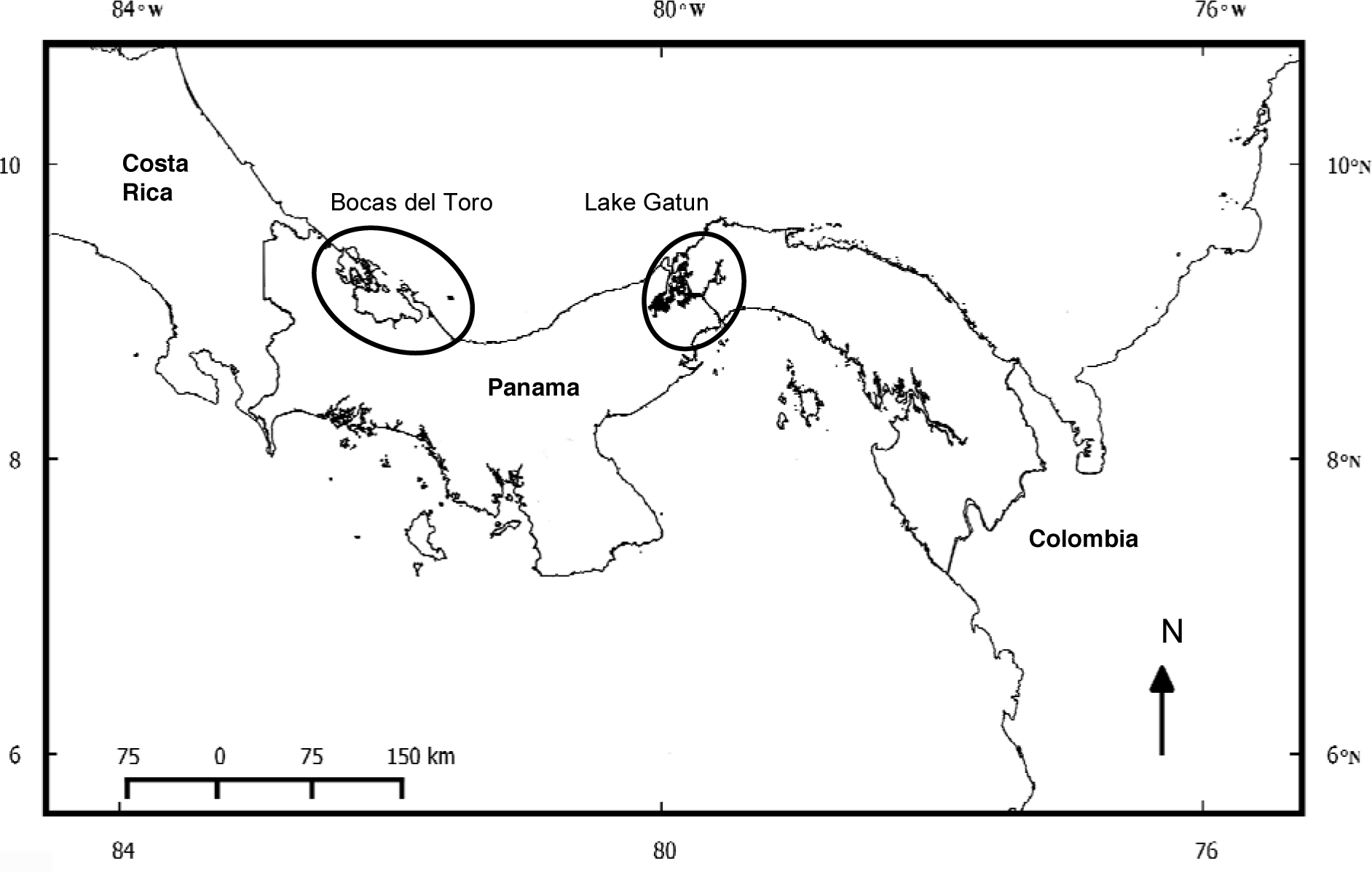
Location of the two known resident populations of West Indian manatees (*Trichechus manatus*) in Panama; Bocas del Toro and Lake Gatun.

The net result of these circumstances is that there is a pressing need to gather information on the current state of this population. In this study we provide preliminary information on the number of the West Indian manatees in Lake Gatun and document possible human threats to manatees in the main body of water within the Panama Canal. Located in central Panama (09°09’N 75°51’W) and covering 430km^2^ (Fig. 2), Gatun is an artificial freshwater lake created in 1906 when the Chagres River was impounded to build the Panama Canal. The typical vegetation in the lake includes *Eichhornia crassipes*, *Pistia stratiotes*, *Pontederia rotundifolia* and *Hydrilla verticillata*, all of which are part of the manatee’s diet (Jimenez-Perez 2000, TLBG *et al*. 2002).

**Figure 2.**
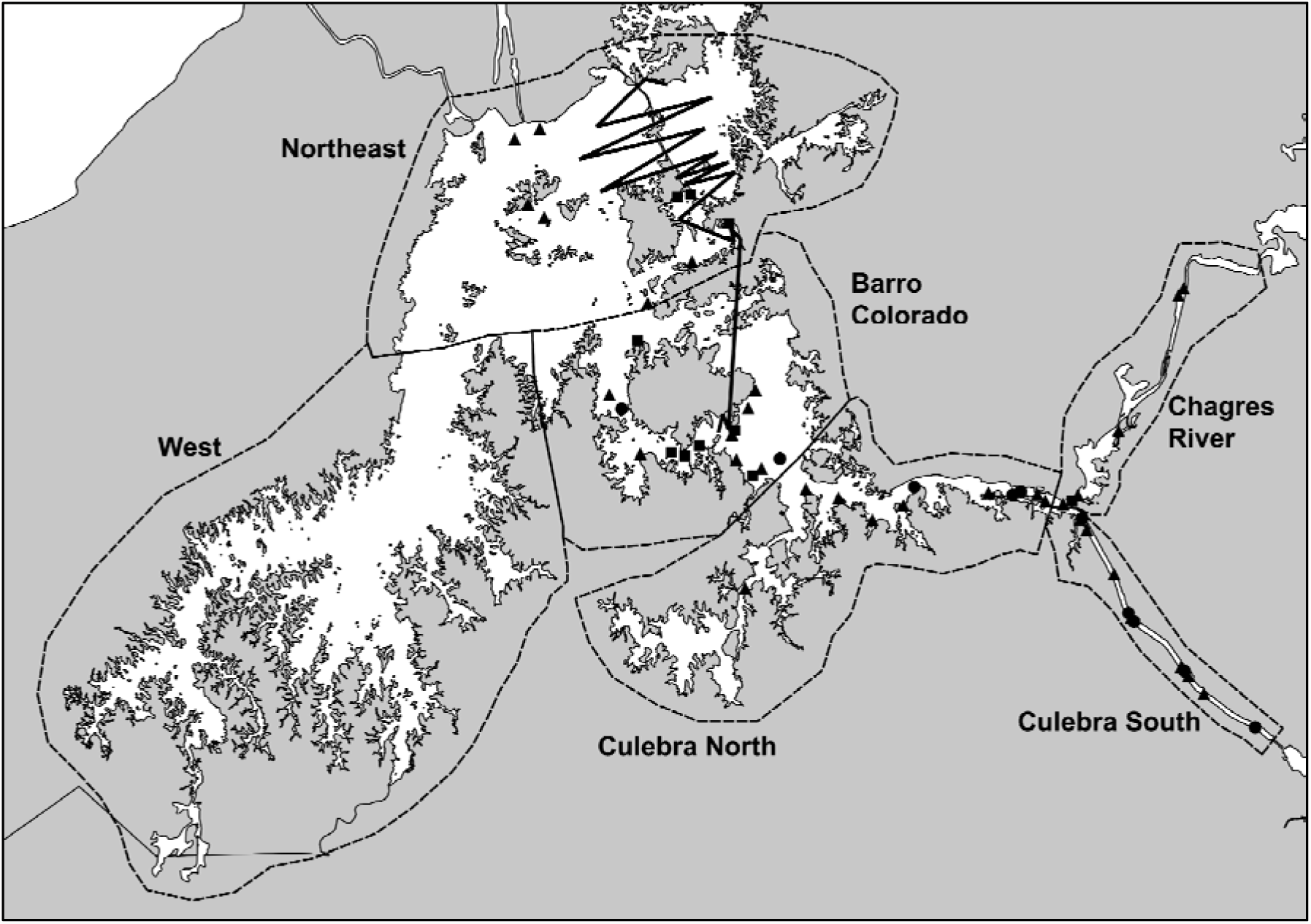
Lake Gatun was divided into six sectors for this study; Sector I Culebra South, Sector II Culebra North, Sector III Chagres River, Sector IV Barro Colorado, Sector V Northeast, Sector VI West. Triangles indicate manatee sightings through interviews, circles indicate sightings from official documents, and squares indicate sightings from boat-based and aerial surveys. Black line indicates transects flown during aerial survey, 10 October 2008.

We conducted interviews with local people in order to determine their perception of and association with manatees. Interview questionnaires were modified from Lima (1997) and Luna (2001) (see Annex 1). Similar to the methodology used by Motonya-Ospina *et al*. (2001), we also reviewed information from the captain’s logs of the Panama Canal Authority (ACP) Aquatic Vegetation Control Unit and the Smithsonian Tropical Research Institute (STRI) Game Warden reports which contained detailed information on the removal of deceased manatees and allowed us to assess the number of recorded manatee carcasses sighted in the lake. Finally, we conducted aquatic (boat-based) surveys and aerial surveys to locate and assess the numbers of manatees in Lake Gatun. We plotted each manatee sighting from each of the different sources on a map using ArcView GIS 3.2 (ESRI, Inc. 2002). We analyzed the proportion of interviews that yielded manatee sightings as well as any grouping in the location of sightings. To simplify analyses, we divided the lake into six sectors: I) Culebra Cut South II) Culebra Cut North III) Chagres River, IV) Barro Colorado, V) Northeast and VI) West (Fig. 2).

We conducted interviews between March and July 2007. Interviews focused on employees of the Gamboa Rainforest Resort, Barro Colorado Nature Monument game wardens, private boat operators, and several Divisions of the ACP since they spend the majority of their working hours (day and night) on the lake. We were not able to conduct interviews with members of local agricultural communities located on the Western sector of the lake due to difficult access. Forty-four interviews were conducted between March and June 2007. Interviewees ranged in age from 27 to 56 years, with a mean of 42 years. Only two of the individuals interviewed had never seen a manatee, and thus the effective number of interviews was 42. These 42 interviews yielded 59 manatee sightings; 63% (37) corresponded to recent manatee sightings (< 3 years), and 22% (13) were historical sightings. Sightings concentrated on Culebra Cut North, Sector II (26%) and Barro Colorado, Sector IV (24%) (Fig. 2). We were not able to conduct interviews with members of local agricultural communities located on the Western sector of the lake due to difficult access.

Only one respondent admitted to hunting a manatee, while 21% (9) of respondents knew of the existing legislation prohibiting hunting of manatees. Of these 78% (6) of which corresponded to the STRI game wardens. 21% (9) recalled seeing females with calves, but there was no specific time of the year when respondents saw calves more often. Interviewees also commonly saw manatees alone (42%) or in pairs (36%); while only 5% reported seeing groups of three or more. Finally, 64% (27) of respondents had seen at least one dead manatee in the lake; 26% (7) speculated they deaths were the result of detonations during dredging activities in Culebra Cut, while 37% (10) attributed deaths to collisions with boats.

A total of 32 Captain’s logs was reviewed, which recorded 19 manatee deaths over the 14 years from 1995 to 2008. The largest number of deaths in one year (n = 4) was registered in 2007. No deaths were recorded during 1998 or 2003. No deaths were reported in February, August, or September of any year. Thirteen deaths (76%) were registered in Culebra Cut (Fig. 2), but the Aquatic Vegetation Control Unit only surveys Culebra Cut, Chagres River and the Tabernilla region (Sectors I, II and IV), so these results must be treated with caution.

Boat-based surveys were carried out 3m long fiberglass boat with a 25hp outboard engine on five different occasions from January to June 2007. Due to difficult access boat-based surveys were only carried out in Sectors II, III and IV. Only two manatees were seen from the boat during the surveys: one in the Chagres River (Sector III) and west of Barro Colorado Island (Sector IV) (Fig. 2).

Aerial surveys were carried out in a Robinson R44 helicopter, flying at an average altitude of 150m and speed of 180km/h (following Lefebvre *et al*. 1995). Restrictions regarding aerial space over the Panama Canal locks required these areas to be flown above 200m. Each flight originated in Marcos A. Gelabert National Airport, Panama City (8.9733° N, 79.5556° W). Aerial surveys were conducted on 10 January 2008, 14 January 2008 and 10 October 2008 (Table 1). All flights were carried out in the early morning to reduce the effect of glare (Marsh and Sinclair 1989). For the aerial surveys Sectors I to VI were flown on 10 January, Sectors I, II, III and V were flown on 14 January 2008, and sectors IV and V on 10 October 2008. The beginning and end of each transect was marked using a GPS (Fig. 2). During each flight, each time a manatee was spotted from the air an observer recorded the transect number, number of individuals seen, if they were in a group (a visually distinct “clumping” of individuals) and the presence of calves (individuals of approximately less than 2m in close proximity to or accompanying a larger individual) (Morales-Vela *et al*. 2000).

**Table 1.** Weather conditions, water clarity, duration and number of manatees sighted during the three aerial surveys, Lake Gatun, Panama Canal, Panama.

| Date | Weather Conditions | Water clarity | Duration of survey | No. of manatees | No. of calves | No. of groups |
| --- | --- | --- | --- | --- | --- | --- |
| 10 Jan 2007 | Good* | Clear | 1.5 h | 10 | 2 | 1 |
| 14 Jan 2007 | Poor* | Clear | 45 min | 2 | - | 1 |
| 10 Oct 2008 | Fair* | Turbid | 1.1 h | 16 | 4 | 2 |

\*as defined by Ackerman (1995) and Hodgson *et al.* (2007).

During the 10 January 2008 aerial survey, 10 manatees (8 adults + 2 calves) were sighted in 1:30 h of survey time. During the 14 January 2008 survey only two manatees (both adults) were sighted in 45 min. During 10 October 2008, 16 manatees (12 adults + 4 calves) were sighted in 1:10hrs. Sightings were made in the Northeast (Sector V) and Barro Colorado (Sector IV) sectors (Fig. 2). There were no sightings of manatees in the Western sector (Sector VI), or either sections of Culebra Cut (Sectors I and II).

Our study, while preliminary, is the first to assess the distribution and conservation status of the West Indian manatee population in Panama Canal in over 20 years. Our results show that are at least 16 individuals in Lake Gatun and that at least some of them are reproducing as evidenced by the sighting of females with young. While 16 manatees is still very few, it is more than what has previously been reported (Mou-Sue *et al*., 1990). Our study also shows the first evidence of manatees reproducing in the Panama Canal.

The 59 manatee sightings obtained through interviews are similar to those obtained by Schad *et al*. (1981). However, we cannot rule out that some sightings were of the same individual and may merely be evidence of a small number of manatees using habitat near areas frequented by humans. For example, Reports in Culebra Cut (Sectors I and accounted for more than 40% of the manatee sightings. Culebra is arguably the busiest section of the Panama Canal. A mere 152m across, the Cut is bustling with activity 24 hours a day, not only due to large shipping vessels (up to 45 a day) but the almost non-stop dredging activity, as well as tourism operators. Manatees could be using the upper portion of Culebra cut to transit to and from the Chagres river and the lake proper. As such, these sightings do not necessarily indicate the level of habitat use or a higher than expected number of manatees in the area. Indeed, there was no apparent correlation between areas of low boat traffic and high sightings of manatees with any of the methods used in this study.

Interviews with residents in the less congested Western sector of the lake would help off-sett this potential bias, but we were unable to interview local farmers and fishermen in this sector of the lake due to difficult access. Past studies have faced a similar challenge (Schad *et al*. 1981, Mou-Sue *et al*. 1990). Future studies would do well to include local communities in this area.

An apparent lack of hunting is encouraging, and is contrary to what has been found in other parts of Panama (Mou-Sue *et al*. 1990), and in the remainder of Central and South America (Reynolds *et al*. 1995, Auil 1998, Luna 2001, Montoya-Ospina 2001, Jimenez 2002). However, it might be that manatees are not yet abundant enough to be a regular source of protein for locals. Results of interviews from communities in the western sector of the lake would go a long way to determining whether or not this is indeed the case. In contrast to studies carried out in Guyana, Brazil and Costa Rica (Reynolds *et al*. 1995, Jimenez-Perez 1998, Luna 2001, de Thoisy *et al*. 2003) few respondents knew of the legislation that prohibits the hunting of manatees (Legislative Assembly 1995). This lack of knowledge needs to be addressed and the legislation needs to be made known to the public at large if manatees are to be protected in the long term.

The 19 reported manatee mortalities from 1995 to 2008 have no accurate cause of death however, manatee mortality in the lake has previously been attributed to collisions with trans-oceanic vessels or under water detonations due to dredging activities in the Canal (Schad *et al*. 1980, Hernández 1982). What the deaths recorded in the captain’s logs and game warden’s reports do reveal that manatees have been consistently present in the lake during those 14 years. A death rate of approximately one manatee per year may not seem high, but considering the small number of manatees in the lake it is cause for concern. Our study represents the first time this type of methodology was used to assess manatee mortality in Panama.

Contrary to what has been found in other studies (LaCommare *et al*. 2012), we only sighted two manatees during boat-based surveys. This is discouraging, as boat-based surveys have proven to be a cost-effective method of monitoring manatee populations in other areas of Central America (LaCommare *et al*. 2012). However, boat-based surveys work best in clear waters and would be of lesser value in the deeper and more turbid waters of Lake Gatun, particularly during the rainy season.

In our study, the highest number of manatees sighted during a single survey, and thus least likely to count individuals twice, was 16. When we consider that aerial surveys tend to underestimate the number of manatees present (Marsh and Sinclair 1989, Lefebvre *et al*. 1995, Edwards *et al*. 2007), and that three aerial surveys are of limited utility in such a large area with dispersed manatees, there may be upwards of 20 – 25 individuals in the lake. Of course, this is still a very small number. By comparison, in Belize and Mexico counts can reach more than 200 individuals (Morales-Vela *et al*. 2000). More locally, in Bocas del Toro Mou-Sue *et al.* (1990) sighted a maximum of 70 manatees, while in Tampa Bay, Florida a single survey can yield 192 sightings (Wright *et al*. 2002) and more recent counts report close to 2000 manatees (see http://myfwc.com/research/manatee/research/population-monitoring/synoptic-surveys/).

However, both Vianna *et al*. (2006) and Tucker *et al*. (2012) found low genetic diversity in the endangered Florida manatee. Their findings suggest that a large population size should not be taken as the main indicator of overall health. In fact, as a “bridge” between Central and South America the preservation of Panama manatees could prove to be relevant to the overall genetic health of manatees in the region (Castelblanco-Martinez *et al*. 2012). We recommend future studies consider assessing the genetic variability of manatees in Panama.

While we did not see any fishing with nets or poles, we did see large expanses of cattle ranching, agricultural and forest crops were clearly visible amongst the small communities that border the Western sector. It is tempting to speculate that the almost permanent human presence at the lake edge may be altering manatee distribution as suggested by Castelblanco-Martinez *et al*. (2012). However, the relatively common sightings of manatees in Culebra Cut could be an indication that at least some manatees are accustomed to human presence and substantial activity. Indeed, the manatee sighted during the complimentary aerial survey of the lake by Mou-Sue et al. (1990) was in this western sector.

The increased agricultural activity in that sector did not alter water clarity during aerial surveys, but turbidity was a factor during the October 2008 survey (Table 1). Water turbidity due to run-off from agricultural lands carried into the lake could have affected visibility. In addition to the Chagres some 20 smaller rivers empty into the lake, which make the water in the lake turbid during the rainy season (from April to November). Reynolds *et al*. (1995) found that in Costa Rica aerial surveys during the rainy season are not a productive way to survey manatees. Turbidity undoubtedly limits the visibility of manatees from the air (Marsh and Sinclair 1989, Lefebvre *et al*. 1995, Edwards *et al*. 2007). Marsh and Sinclair (1989), Lefebvre *et al*. (1995) and Edwards *et al*. (2007) discussed the problems of aerial surveys for the estimation of population sizes and trends. These results highlight the importance of using a variety of methods to assess population status of cryptic species (Hines *et al*. 2008). We recommend not only more intense aerial surveys in the future, but also assessments using side-scan sonar which has been proven highly effective to detect manatees, particularly in turbid waters (Gonzalez-Socoloske *et al*. 2009).

In conclusion, the number of manatees present in the Panama Canal, while still quite small, is slightly higher than previously thought and we now have evidence that they are reproducing in the lake. However, there is a relative high number of deaths which is cause for concern. While hunting does not appear to be an immediate threat, the impact of large transoceanic vessels and underwater detonations for dredging on the manatees in lake Gatun would need to be assessed. A manatee carcasses recovery program needs to be instated in order to perform proper necropsies to determine accurate causes of death and to recover biological samples from the deceased animals, such as the protocol described by Bonde *et al*. (1983, 2012). We also recommend further, more detailed surveys in order to understand population trends and habitat use.

## Acknowledgements

This work was funded by Wildlife Trust Alliance and The Rufford Foundation’s Small Grants for Nature Conservation. We wish to thank the Aquatic Vegetation Control Unit of the ACP, the Gamboa Rainforest Resort, STRI and the BCI Game Wardens. Special thanks to Isis Tejada, Karla Aparicio, Daniel Muschett, Milton Clark, Ana Salazar, Apolonio Vásquez, Oris Acevedo, Belkys Jimenez and Narkis Morales for their invaluable assistance.

**Annex 1** Sample of survey questions (translated from Spanish) applied to personnel of the Gamboa Game Forest Resort, the Panama Canal Authority (ACP), the Smithsonian Tropical Institute (STRI), and private boat operators between March and July 2007 to determine their knowledge of manatees in Lake Gatun, Panama Canal. Questionnaire modified from Lima (1997) and Luna (2001).

Date:________Survey number:________

Name:__________________________Age:__________

Occupation:_______________ Organization:_____________

Department:_______________

Location:_______________

Time employed__________(months, years)

Please tick ( ):

Have you ever seen a manatee? ( ) yes ( ) no

Where did you first see one?______________________________

Do you remember when? ( ) yes_______________ ( ) no

Can you describe a manatee?______________________________

Have you seen a manatee more than once? ( ) yes ( ) no

When was the last time you saw a manatee?______________________________

Where have you seen them most often?______________________________

-or mark the locations where you have manatees them on the map provided What was the manatee(s) doing?

( ) feeding ( ) traveling ( ) breathing at the surface ( ) other_______________

Is there a time of the year when manatees are more common?

( ) dry season ( ) rainy season ( ) don’t know

Have you seen more than one manatee together in a group? ( ) yes ( ) no

- what is the largest number of manatees you have seen in a group? largest number (_______________) smallest number (_______________)

Have you seen females with calves? ( ) yes ( ) no

-when did you see them?______________________________

Have you seen a dead manatee? ( ) yes ( ) no

- do you know the cause of death?______________________________

Have you ever hunted a manatee? ( ) yes ( ) no

-how is the hunt carried out?______________________________

Do you know of the laws that prohibit hunting of manatees? ( ) yes ( ) no

